# Optimizing biomedical information retrieval with a keyword frequency-driven Prompt Enhancement Strategy

**DOI:** 10.1101/2024.04.23.590746

**Authors:** Wasim Aftab, Zivkos Apostolou, Karim Bouazoune, Tobias Straub

## Abstract

**Background:** Mining the enormous pool of biomedical literature to extract accurate responses and relevant references is a daunting challenge, made more difficult by the domain’s interdisciplinary nature, specialized jargon, and continuous evolution. Early natural language processing (NLP) approaches faced considerable hurdles in comprehending the nuances of natural language which frequently led to false answers. However, the discovery of transformer models has immensely boosted the cause by enabling researchers to create larger and more potent language models known as large language models (LLMs). These LLMs have opened up new possibilities for enhancing question-answering (QA) tasks. Even with these technological advances, the current LLM-based solutions for querying specialized domains like biology and biomedicine still have issues in generating up to date responses while preventing “hallucination” or the generation of plausible but factually incorrect responses.

**Results:** This paper focuses on prompt enhancement using retrieval-augmented architecture as a strategy to guide LLMs towards generating more meaningful responses for biomedical question-answering tasks. We evaluated two prompt enhancement approaches using GPT-3 and GPT-4, examining their effectiveness in retrieving relevant information from complex landscape of biomedical literature. Our proposed approach leverages explicit signals in user queries to extract meaningful contexts from a vast pool of information, addressing the shortcomings of traditional text embedding-based prompt enhancement methods. We developed a QA bot ‘WeiseEule’ (https://github.com/wasimaftab/WeiseEule-LocalHost) utilizing these prompt enhancement methods and allowing for comparative analysis of their performances.

**Conclusions:** Our findings highlight the importance of prompt enhancement methods that utilize explicit signals in user’s query over traditional text embedding based counterparts to improve LLM-generated responses in specialized domains such as biology and biomedicine. By providing users complete control over the information that goes into the LLM, our approach tackles some of the major drawbacks of existing web-based chatbots and LLM-based QA systems including hallucinations and the generation of irrelevant or outdated responses.

## Background

Retrieving precise answers and relevant references from large volumes of text is still a crucial but challenging task in biomedical research. The complexity of this endeavor arises due to several unique characteristics and challenges inherent to biomedical literature. The language used is highly specialized and includes a lot of acronyms, technical jargon, and complicated medical terms. Moreover, the biomedical domain is ever evolving, featuring constant improvements and breakthroughs. Because of this rapid advancement, new information is constantly being added, making it difficult for the retrieval systems to stay updated and in line with the most recent advancements. Furthermore, biomedical information is shared in various formats such as research papers, clinical trial reports, patent documents and so on. The structural complexities of each of these formats make it difficult to extract and interpret information with accuracy.

In addition to these challenges, biomedical literature uses synonyms, and highly context-dependent phrases which adds yet another level of complexity. Because language usage varies significantly, ambiguities may arise that make it challenging for information retrieval systems to understand the precise context or meaning intended in a query. Another thing is that biomedical research often intersects with other disciplines such as statistics, physics, and computer science etc. Therefore, to deliver thorough and pertinent information, retrieval systems must be able to integrate and understand concepts from these many domains.

Early NLP approaches which primarily used a mixture of rule-based algorithms, statistical analysis, and classical machine learning techniques made progress in solving these issues [1–4]. However, their ability to comprehend subtleties and context of complex biomedical texts was often limited, and they struggled to capture long-term dependencies in the text. This frequently resulted in inaccurate or irrelevant responses.

The emergence of transformer models has revolutionized the field of NLP. A much richer understanding of context and linguistic nuances is made possible by Transformers, which process entire text sequences simultaneously by utilizing deep learning and a novel use of self-attention mechanisms to process sequences of text [5]. Transformer based models have demonstrated potential in capturing long-term dependencies in text, better interpreting context, and comprehending the nuances of natural language, resulting in improved accuracy and relevance in generated responses compared to earlier models [6–8].

The development of LLMs have further enhanced QA tasks, building on the revolutionary developments initiated by early transformer models [6–9]. LLMs are instances of foundational models that are pre-trained on large amounts of unlabeled and self-supervised data, meaning the model learns from patterns in the data to produce generalizable and adaptable output. And LLMs are those instances of foundational models that are applied specifically to text and text like entities such as codes. These models can be tens of gigabytes in size and trained on enormous amounts of text. For instance, GPT-3 from OpenAI, is pre-trained on a corpus of 45 terabytes of text corpora from the web potentially encompassing scientific and biomedical literature [8]. In the context of biomedical QA, LLMs can generate answers to complex questions by leveraging their extensive pre-training and with an increase in the number of parameters, they typically demonstrate enhanced performance [8, 10]. These parameters essentially represent the size of an LLM’s “brain,” and we are living in an era where we have a competitive trend in creating LLMs with ever bigger brains. Among many available LLMs, OpenAI’s GPT-3 and 4 have gathered significant attention after their chatbot ChatGPT has gone viral globally [11]. ChatGPT offers a browser interface for easy access to OpenAI’s LLMs, but these models can also be accessed via Application Programming Interfaces (APIs) for more customized applications such as querying a large pool of personalized information. However, training LLMs daily with newly added information is prohibitively expensive therefore, they suffer from a major issue: possibility of yielding outdated responses due to not keeping up with current literature. In addition, LLMs suffer from hallucination which in the current context refers to the phenomenon where the model generates responses that may sound plausible but are not factually correct. While OpenAI has integrated Bing search capabilities for ChatGPT plus subscribers to closely align responses with the currently available information on the web, this feature is presently limited to web-based access only, eventually limiting its applicability to nontrivial NLP tasks such as biomedical QA. Moreover, even with dynamic web access, ChatGPT plus users are limited to the initial search results otherwise speed becomes a bottleneck and in fact for many complex queries, ChatGPT with Bing search takes longer time to generate answers.

Another player in the web-based conversational search engine space is Perplexity.ai which aims to generate answers using the LLMs such as GPT-3.5, 4 and Claude, Mistral Large, and an in-house model which is currently experimental [12]. Though Perplexity.ai supports querying custom knowledge base via file uploads for premium users, it has restrictions on the number and size of the files. Therefore, the knowledge base of these chatbots that aim to cater all domains, are substantially limited to the information that is available freely on the web. This limitation can significantly affect the quality of responses from QA systems, especially in rapidly evolving fields like biology and biomedicine.

Researchers affiliated to institutes and universities typically have access to vast amounts of research content via institutional subscriptions. Additionally, they often obtain research papers through networks of fellow researchers in other universities, accumulating extensive personal libraries over time. Querying this huge pool of literature is beyond the capabilities of current web-based chatbots, indicating a gap in the applicability of these tools for thorough question answering.

More recently, researchers have proposed prompt engineering (PE) approaches for variety of use cases to improve responses generated by an LLM [13–15]. A “prompt” refers to a textual input provided by the user to instructs and/or guides an LLM to generate meaningful responses and prompt engineering is mainly about providing as much context as possible with providing few examples (few-shot learning) to enhance an LLM’s understanding of a given use-case. However, PE leverages pre-trained knowledge base wired into the parameter space of an LLM which can lead to issues like outdated or fabricated responses mentioned earlier. Therefore, its effectiveness is somewhat limited in certain applications such as question answering (QA) especially within specialized domains like biomedicine. To address these issues, researchers have proposed prompt enhancement strategies (PESs) utilize a retrieval-augmented generation (RAG) based architecture which aims to dynamically extract contexts from a vast pool of information to augment user’s prompt [16]. This augmented/enhanced prompt then serves as the knowledge base for an LLM to seek answers.

In this paper, we focus on improving information retrieval for biomedical QA tasks by utilizing RAG based PES. Within open domains QA tasks, PES based on retrieval-augmented architecture is quite popular approach. It mainly relies on text embedding and similarity of vectors in a high dimensional space to extract contexts for prompts [17–19]. While this PES provided substantial progress toward addressing the issues with LLMs, we found it fall short, particularly when dealing with specialized biomedical questions. Therefore, there is a need for a strategy that addresses the shortcomings of text embedding based RAG by more effectively navigating the complex landscape of biomedical literature. To address this issue, we developed a prompt enhancement method based on retrieval-augmented architecture but instead of using similarity of vectors it uses explicit signals from user queries to extract relevant contexts from the vast pool of biomedical literature. Interested users can compare the performances of the approaches discussed here in a QA bot called ‘WeiseEule’ (See Availability of data and materials).

## Methods

The conceptual underpinning of our algorithm is depicted in Fig 1. The idea is to first create a knowledge base from research papers. Then using prompt enhancement techniques based on retrieval-augmented architecture (see sub section 3,4) construct relevant prompts that can be sent to LLM for generating answers to user questions. In the following sub sections, we describe the details of the prompt enhancement approaches.

**Fig. 1:**
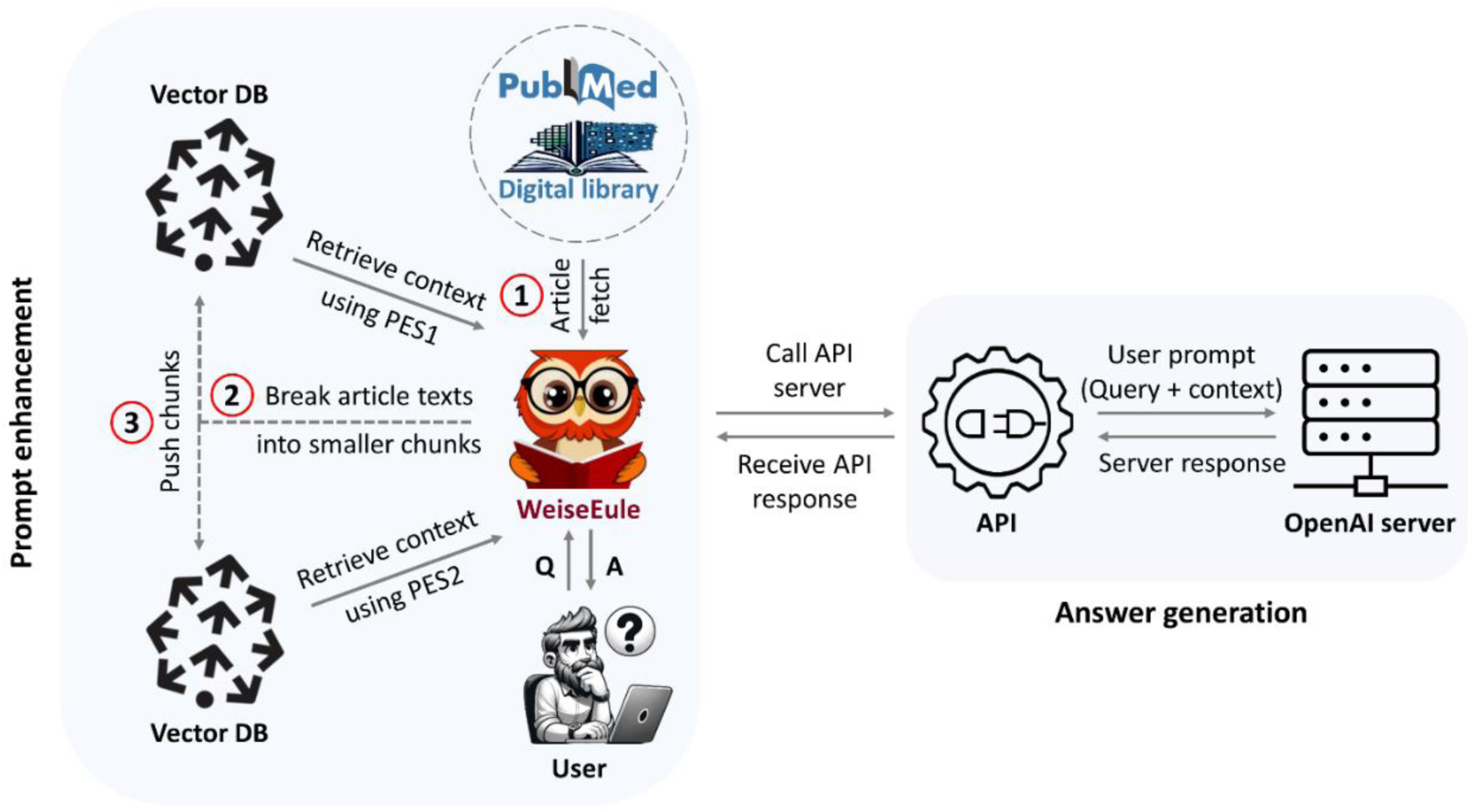
Framework of the WeiseEule pipeline –. The WeiseEule pipeline is structured to facilitate a seamless QA experience, combining namespace creation, and generating capacity of LLM with strategic prompt design. Three necessary steps required to set up the knowledge base is indicated in red circles. Users can choose between PES1 and PES2 approaches to retrieve contexts.

### 1. Knowledge base

The first and the most crucial step is to generate a knowledge base (KB), which helps users to have total control of information that goes into the LLM and massively reduces the chances of hallucinations. To build a custom KB one can download articles from PubMed in text format or can even utilize already accumulated PDFs in a personal digital library. However, the most important task here is to break the article text into smaller pieces (chunks) so that they can fit into a prompt. Also, breaking down text into smaller chunks enable LLMs to focus on smaller, contextually meaningful units rather than complete documents, which can enhance performance. Then each of these chunks need to be converted into sequence of numbers so that they can be used during text embedding based PES described later. Those vectors are subsequently pushed into a vector database (DB) for high performant access. Key to our KB creation is the concept of namespace (see Figure 2). A Namespace serves as a container to hold relevant information in place. Say for example, a user has a query about *dosage compensation* which refers to the mechanism organisms use to balance gene expression across different biological sexes. Most likely the answer will be found in papers that have *dosage compensation* listed as keyword. So, if one accumulates all those papers and put them in a virtual container, then it forms a namespace, provided it also receives a name tag, say “dosage compensation papers”. Thus, a namespace with the chunks and vectors serves as a custom knowledge base for the prompt enhancement strategies discussed in this paper. In a vector DB, there can be multiple namespaces generated from different topics and namespacing approach helps in organizing heterogeneous knowledge bases in a single vector DB or in one cloud account. However, for namespacing to function correctly, the initial step is to convert the text chunks within a namespace into vectors as described in the next section.

**Fig. 2:**
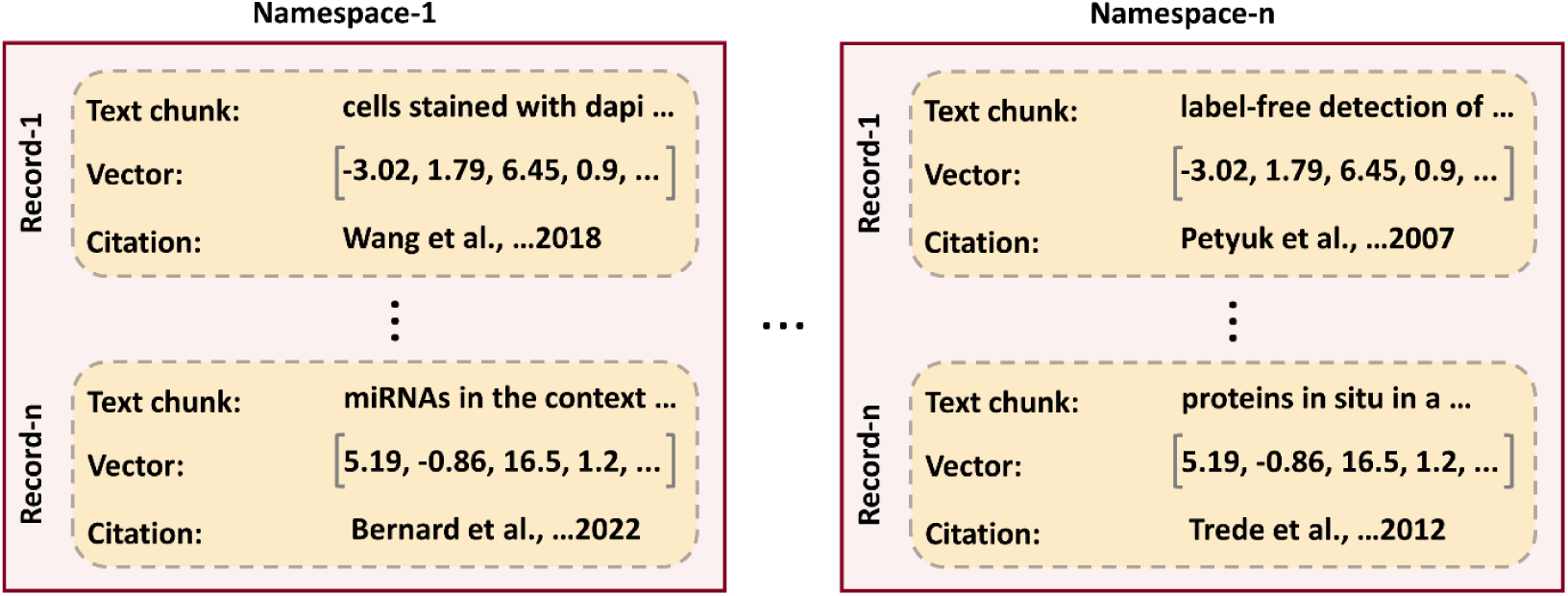
Namespacing –. The concept of namespacing is foundational to our knowledge management approach. Within vector databases, namespaces serve as virtual containers that store relevant information. Each record in a namespace is a composite data structure that can hold a variety of metadata, such as the text chunk, its corresponding vector representation, and bibliographic references to its original source.

### 2. Text to numbers

The transformation of text into embedding vectors is called text embedding. There are specialized models, called *text embedders*, responsible for this [6, 20, 21]. This conversion retains the text’s semantic meaning (Fig. 3), ensuring the LLM captures relationships between different text parts. Text embeddings are a type of representation that allows words with similar meanings to be represented similarly (Fig. 3). It is a projection of text into a high dimensional latent space resulting in compression of text into a list of floating-point numbers (vector). This is typically done by learning dense/compressed representation through context. In this study, we compared three different text embedders:

- <u>text-embedding-ada-002</u>: It is an improved version of previous generations of embedding models such as text-search-davinci-*-001 and text-search-curie-*-001 etc. from OpenAI [22]. This model was trained on large amount of data form general domain texts such as Wikipedia [23].
- <u>BioBERT-v1.1</u>: This model was obtained after pre-training the Bidirectional Encoder

**Fig. 3:**
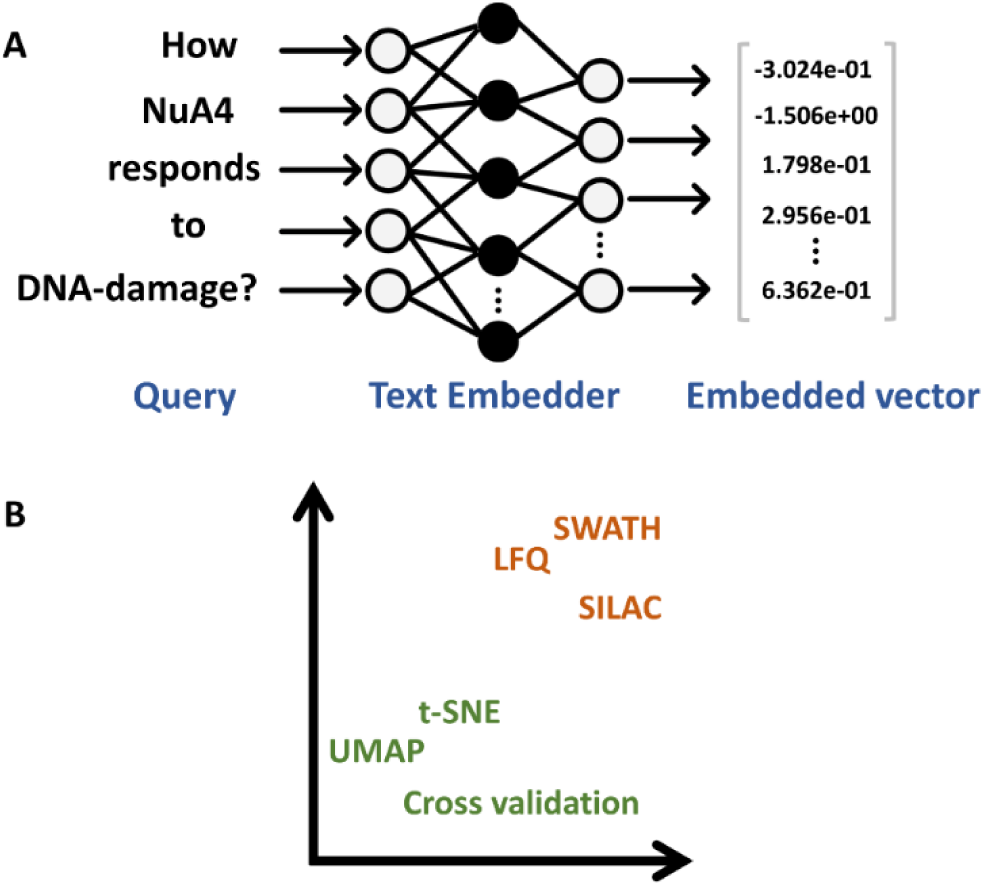
How an LLM understand texts –. An LLM interprets texts by first turning them into embedding vectors. **(A)** The conversion aims to preserve the semantic meaning of the text by making the LLM to capture as much as possible the connections between various text parts. **(B)** Text embeddings are a sort representation that allows words with similar meanings to be represented similarly.

Representations from Transformers (BERT) language model on large amount biomedical text [24].

- <u>BioGPT</u>: This model was obtained after pre-training Generative Pre-trained Transformer

(GPT) model on massive number of abstracts and titles from PubMed articles [25]. The pre-training was performed on GPT-2 model architecture.

Once the chunks are converted into vectors, they need to be stored in a vector DB for faster and efficient querying. The aim is to efficiently extract chunks relevant to construct high quality prompt for a given query. This approach forms the core of the prompt enhancement approach discussed in the next section.

### 3. Context retrieval based on vector similarity ranking (PES1)

To store the vectors, we choose Pinecone DB because it provides ultra-low query latency even with billions of items [26]. When a user asks a question, it is converted to a vector which is then searched in vector DB to extract chunks that have high similarity with the query vector. The cosine similarity (*S_θ_*) between a query vector (q) and the vector corresponding to some chunk (c) is given by equation (1)

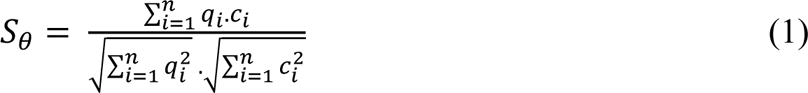

The main reasons to use cosine similarity as a metric to compute similarity between vectors is as follows:

a. <u>Semantic similarity</u>: It measures the cosine of the angle between the two embedding vectors which effectively captures the semantic similarity between them [27].
b. <u>Normalization</u>: It considers the direction of the vectors rather than their magnitude by normalizing for the length of the vectors. This is especially helpful in text mining applications where though the frequency (and thus the magnitude) of words can vary significantly, but it’s the directional similarity that conveys meaningful semantic relationships [28].

In vector DB, every vector has a pointer to a corresponding chunk. Thus, by referring to *top k* similar vectors we can actually extract corresponding chunks which is then used to construct user prompt. In this prompt engineering approach, we first extract the *top k* relevant chunks, then wrap them around user query and instruct the LLM to find answer within these chunks only and if the answer is not found then respond with ‘I don’t know’ (see Fig 4A).

**Fig. 4:**
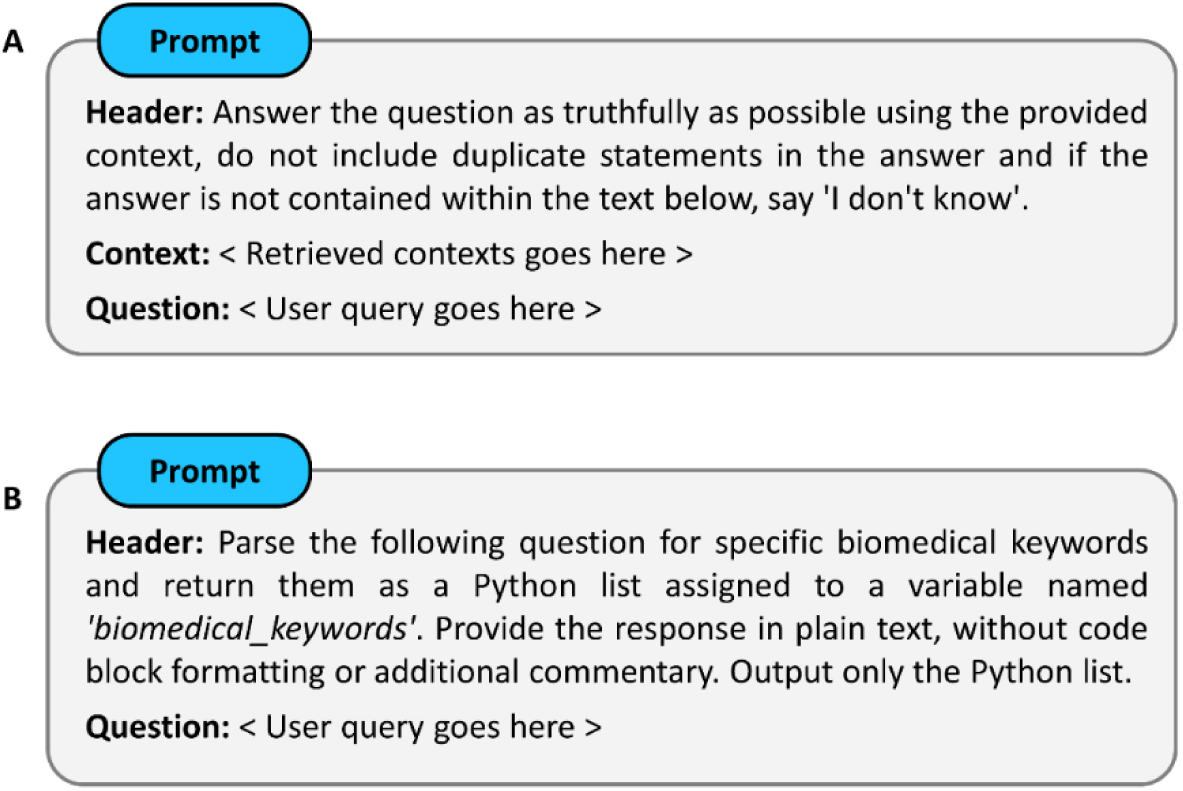
Architecture of our prompt –. Custom instructions in the header guides an LLM to produce desired outcomes. **(A)** Reinforces an LLM to produce improved responses by combining the elements of information retrieval and generative AI. **(B)** Exploits LLM’s zero shot reasoning capability to transform it into an automatic keyword extractor module.

However, it is important to note that this choice of *top k* is very much dependent on the context lengths provided by LLMs (Table 1). Therefore, ideally the top chunks should contain relevant information to provide a satisfactory answer to the question. Yet, this is not always the case (See Results, sub-section 1), which makes the QA bot perform poorly. Therefore, in order to find the most relevant chunks for a given query we developed a prompt enhancement approach that considers the frequencies of keywords as a feature to rank chunks as described in the next section.

**Table 1:** LLMs and their context lengths.

| Model | Context lengths | Max fit |
| --- | --- | --- |
| GPT-3 | 4K tokens ≈ 16K chars | 8 chunks (7 context + 1 answer*) |
| GPT-4 | 8K tokens ≈ 32K chars | 16 chunks (15 context + 1 answer*) |

\*Assuming answer length is ≤ 2000 characters

### 4. Context retrieval based on keyword frequency ranking (PES2)

The aim is to extract keywords from user’s query and rank paragraphs/chunks based on their frequencies. The keyword extraction can be done either manually or automatically. Using well designed and explicit instruction headers within the prompt, it is possible to make an LLM to function as automatic *keyword extractor* (see Fig 4B). Our QA bot implementation offers both options. For automated extraction, LLM outputs the keywords as a python list. However, in some scenarios, it might be necessary to exclude certain keywords or combine adjacent keywords into a single item. In such cases a manual approach is more appropriate. For manual extraction, queries are prefixed with a ‘#’ symbol for ease of programming and keywords are marked using double asterisks. For example, **#**Find all results that connect **NuA4** with **meiosis**.

The approach is based on the consideration that chunks that have a higher frequency of the extracted keywords are more likely to contain useful contextual information to generate a high-quality response. Let us understand this using an example: suppose the topmost table in Fig. 5 is obtained after ordering chunks using cosine similarity-based ranking. If we stick to our considerations, the second row should clearly have come first. To correct that, we re-rank the chunks (Fig. 5). This approach elevates the most relevant chunks by analyzing keyword frequencies, ensuring key information surfaces to the top. By default, and as shown in Fig. 5, we only keep chunks in which all keywords occur at least once. However, several other types of ranking are possible (Fig 6), some of which are described below.

**a.** <u>None fixed</u>: A chunk is ranked higher if it contains all the three keywords over a chunk that does not. Even if the total frequency count is the same or even higher. When two chunks have nonzero values for all keywords, the one with higher ‘total_count’ is ranked higher. In this mode equal importance is given to all the keywords.
**b.** <u>One fixed</u>: More importance is given to chunks that have nonzero values in the fixed keyword column which in this case is ‘k3’. This way of ranking chunks is based on Boolean expressions (k1 OR k2 AND k3), similar to searches in PubMed.
**c.** <u>Multiple-fixed</u>: This is a generalized version of *one fixed* mode. When a user wants to fix multiple keywords, this mode can be useful.

**Fig. 5:**
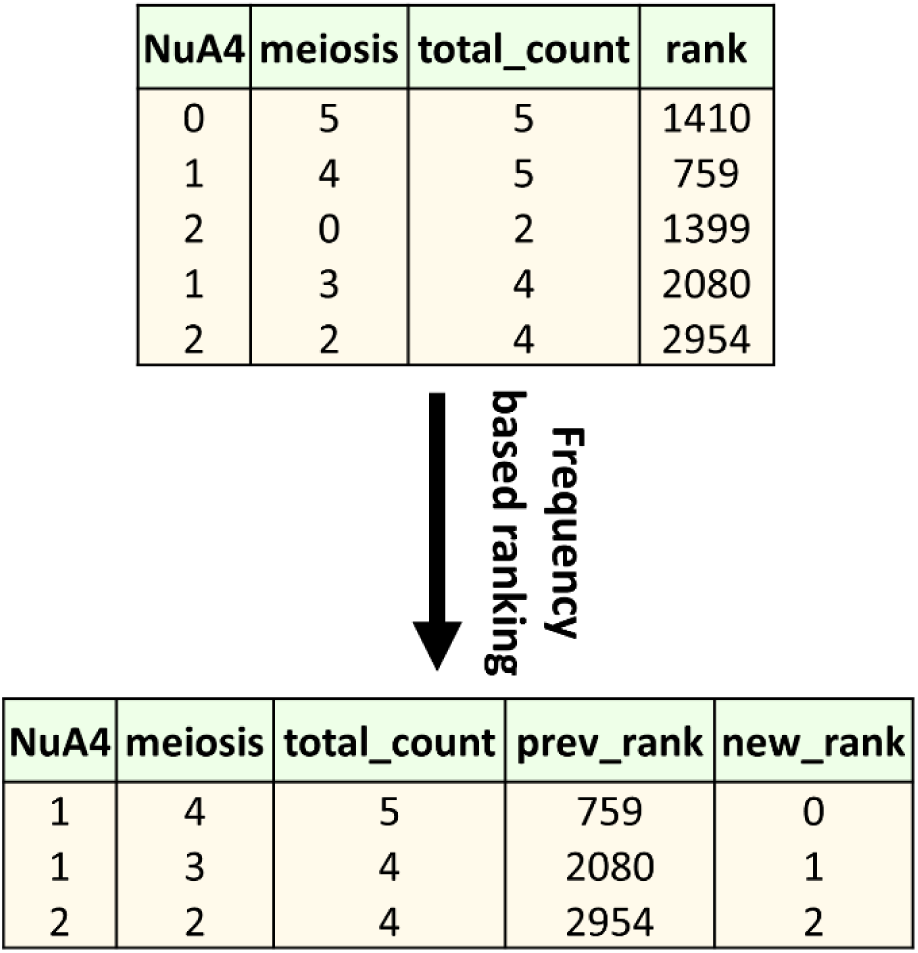
Re-ranking chunks –. ‘NuA4’ and ‘meiosis’ are keywords extracted from user query: ‘Find all results that connect NuA4 with meiosis’. Each row represents a paragraph or chunk from research papers, with columns displaying both individual and total frequencies of the keywords. The upper table shows chunks that are initially ranked according to cosine similarity computed between the query and each chunk in the vector DB. The bottom table depicts the re-ranking of the same chunks based on their keyword frequencies.

**Fig. 6:**
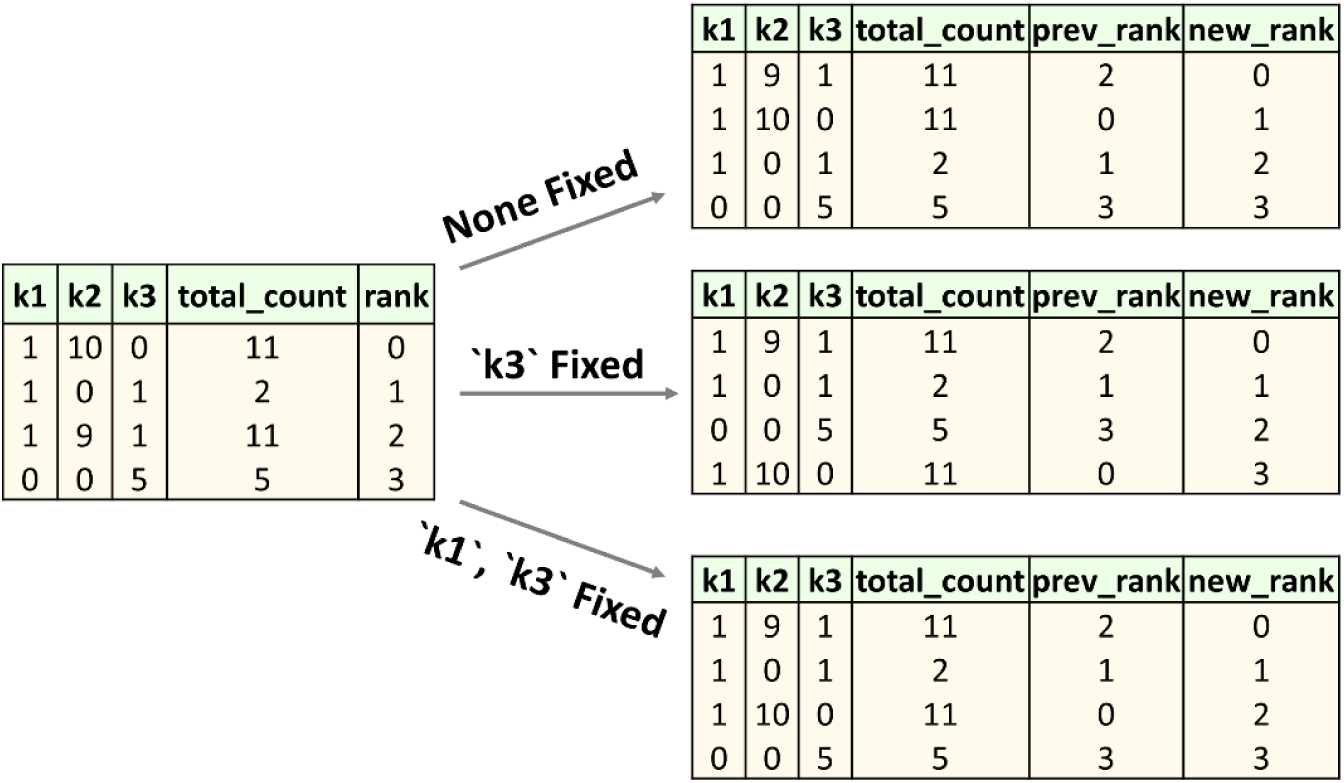
Comparative Chunk Ranking Based on Keyword Frequencies. – Each row represents a text chunk, columns indicate both individual and total frequencies of three keywords: k1, k2 and k3. The left table presents chunks initially ranked according to cosine similarity. The successive tables on the right depict the re-ranking of these chunks under different ranking criteria: without any fixed keywords (None Fixed), with one keyword (‘k3’) is fixed, and with multiple keywords (‘k1’, ‘k3’) fixed, respectively.

Once the chunks have been extracted and a context for answering the user query is created, processing takes place via LLM.

### 5. Processing queries by accessing LLMs via API

The extracted context is wrapped around the user query and a prompt for LLM is created. We use the API services from OpenAI for customized interaction. In the prompt header, the LLM is explicitly instructed to look for the answer only within the given context and if the answer cannot be found then it should return appropriate response, such as ‘I don’t know’ (see Fig 4A). Thus, by instructing the model to search within the provided passages, we give it access to the latest data and benefit from its contextual understanding and ability to generate human-like responses.

Our pipeline (Fig 1) can be divided into two main modules: one for designing relevant prompts and the other for generating meaningful responses. At the heart of the *prompt enhancement* module lies the text embedding, which aims to capture the essence of text in a compressed representation that can be utilized to obtain relevant contexts. At the same time, the core to the *answer generation* module is the LLM responsible for generation human like responses. As users of AI models, we are bound by certain constraints: For instance, creating a proprietary LLM is currently prohibitively expensive for most users. Despite that, we can still influence the outcome by tweaking the other aspects of our pipeline. Our hypothesis is that text embedding has a significant impact on the final response generated by LLM. Because it is through these embeddings and their similarities with the query that we determine the context for our prompts.

### 6. Experiment: Influence of text embedding on answer quality

Before discussing the experiment, it is important to briefly note some of the key aspects of text embedding. Vector embeddings can be seen as a form of compression, a way of capturing the essence of text, i.e., semantic similarity between words or sentences in a format that a machine can understand and manipulate. It has been shown that the dimensionality of embeddings can impact model’s understanding of semantic relationships [21, 27]. There are several text Embedder models available, and they embed the texts into different dimensions. For example, the OpenAI model embeds text into 1536 dimensions. A larger dimension might capture more information, but it also requires more computational resources to process and store. Higher-dimensional vectors might also include dimensions that capture noise rather than useful information, leading to overfitting. Conversely, reducing the dimensionality might lead to information loss, but it can help focusing on the most important aspects of the text’s properties and result in models that are more efficient and less prone to overfitting. Consequently, there is a trade-off which a practitioner needs to optimize. Also, text embeddings are domain sensitive because the meaning of words can differ based on context. For example, “cell” in a biology text refers to a basic unit of life, but in a telecom context, it refers to a cellular network. Therefore, researchers have published several embedding models that are fine-tuned/re-trained on data from a specific domain, such as BioBERT which is BERT model pre-trained on large-scale biomedical texts. BioGPT is a GPT model in its core but again trained on vast amounts of texts from bio literatures. Based on these facts we hypothesized that different text embedder models can influence the quality of answers generated by LLM. We therefore tested which of the three models: BioBERT, BioGPT and text embedder ADA model from OpenAI works best for our knowledge management use case. The experiments were performed as follows: we first created a ‘Knowledge base’ where papers from a specific topic are stored after chunking article texts into small paragraphs of 2000 characters (chars) long (as depicted by the small boxes in Fig 7). This specific chunk size was determined through trial and error. Experimenting with various chunk sizes may be helpful to evaluate how they influence the precision of the generated answers. We then formed a question, already knowing which chunks contain the answer, which we refer to as *key chunks*. Next, we modified our knowledge base to contain only one key chunk (while rests are non-key chunks) and we asked which embedding model can yield vectors corresponding to the key chunk and our query, so that the similarity between them is the highest. In other words, we tested which embedding model can rank the key chunk the highest, keeping in mind LLMs like GPT-3 and GPT-4 can only consider the top 8 and 16 chunks, respectively (see Table 1). This entails that any chunk ranked lower than these thresholds will be missed by the corresponding models.

**Fig. 7:**
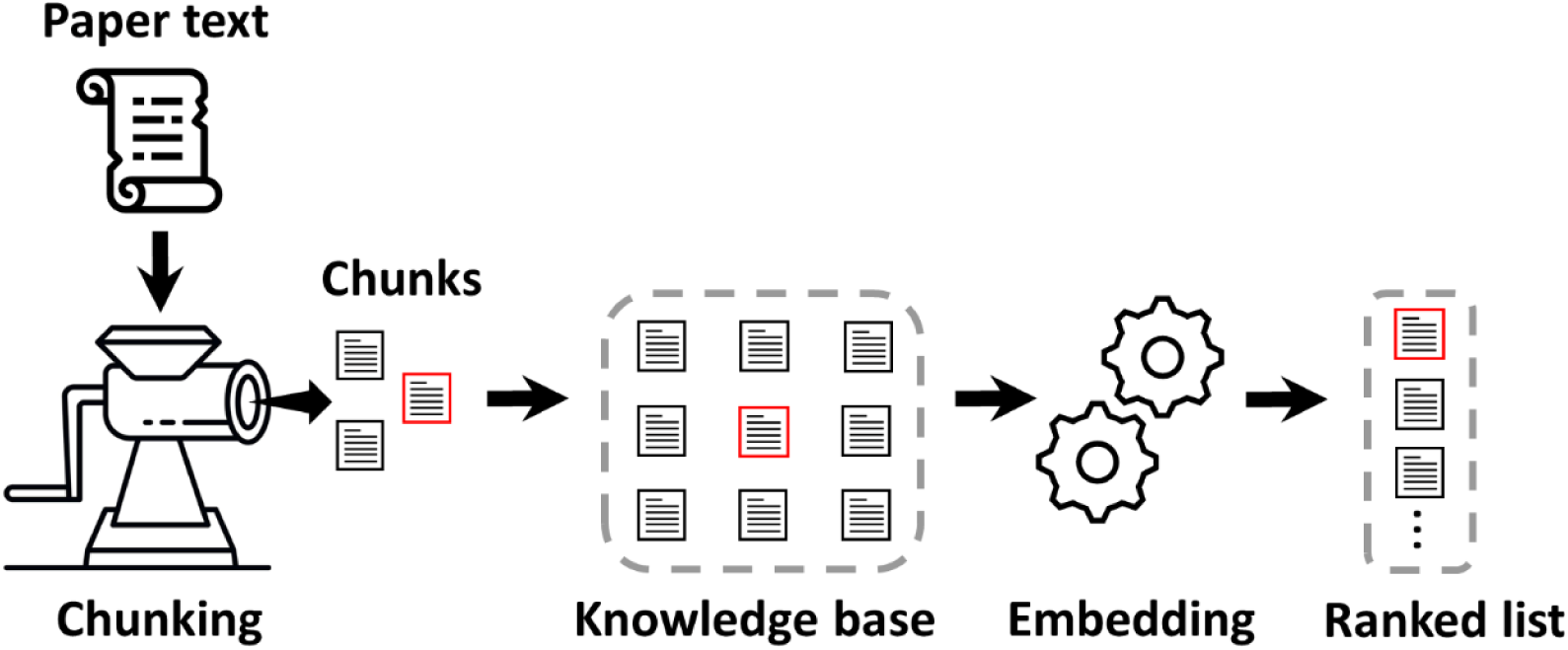
Experiment to determine optimal embedding model –. Text from research papers is broken into smaller chunks to construct a knowledge base. A key chunk (which contains the answer) is highlighted within a red rectangle. The objective is to evaluate which embedding model assigns the highest rank to the key chunk.

## Results

### 1. PES1 struggles with optimal prompt generation in relatively large knowledge bases

In our first setup, the knowledge base contained just 7 papers and one known key chunk. Our query was: *Find all results that connect NuA4 with meiosis*. We found that both BioBERT and BioGPT ranked the key chunk 2^nd^. OpenAI’s ADA, though, ranked it 9^th^. So, if we use ADA with GPT-3’s limit, then the key chunk would be missed and hence the answer

Next, we wanted to further evaluate whether this pattern was consistent with other larger namespaces using the same query as above. Hence, in our second experiment, we selected ≈200 papers and, this time, other chunks could possibly contain the answer as well (which we did not know). BioBERT again outperformed others and placed the key chunk at the 29^th^ position. However, it was beyond the reach of both GPT-3 and GPT-4’s limits. Furthermore, we inspected the chunks ranking above the 29^th^ place and found that they lacked sufficient context to answer the question as effectively as the key chunk. This demonstrate that the choice of embedding has a significant impact on the answer. Observing this, we realized we needed to boost the key chunk’s visibility and we achieved that with PES2.

### 2. PES2 improves prompts by boosting visibility of relevant chunks

We employed PES2 on our knowledge base with about 200 papers, and we used the same query as before. From Fig 8, it is clear that the re-ranking process significantly improved the ranking of the key chunk, lifting it from 29^th^ to 2^nd^ place, surpassing the initial ranking given by BioBERT, the top-performing embedding algorithm from PES1. Interestingly, it also pulled up some additional relevant chunks which were low-ranking before. We subsequently evaluated PES1 and PES2 based on the quality of answers generated by the LLM. A set of ten such questions-answers are provided in Table 2 to illustrate this evaluation. These questions were curated from a diverse range of concepts within chromatin biology; the concepts are color-coded in the table for clearer categorization. These are not so obvious questions and many of them seek extremely specific information. The concepts are segregated in different namespaces as shown in Table 3.

**Fig. 8:**
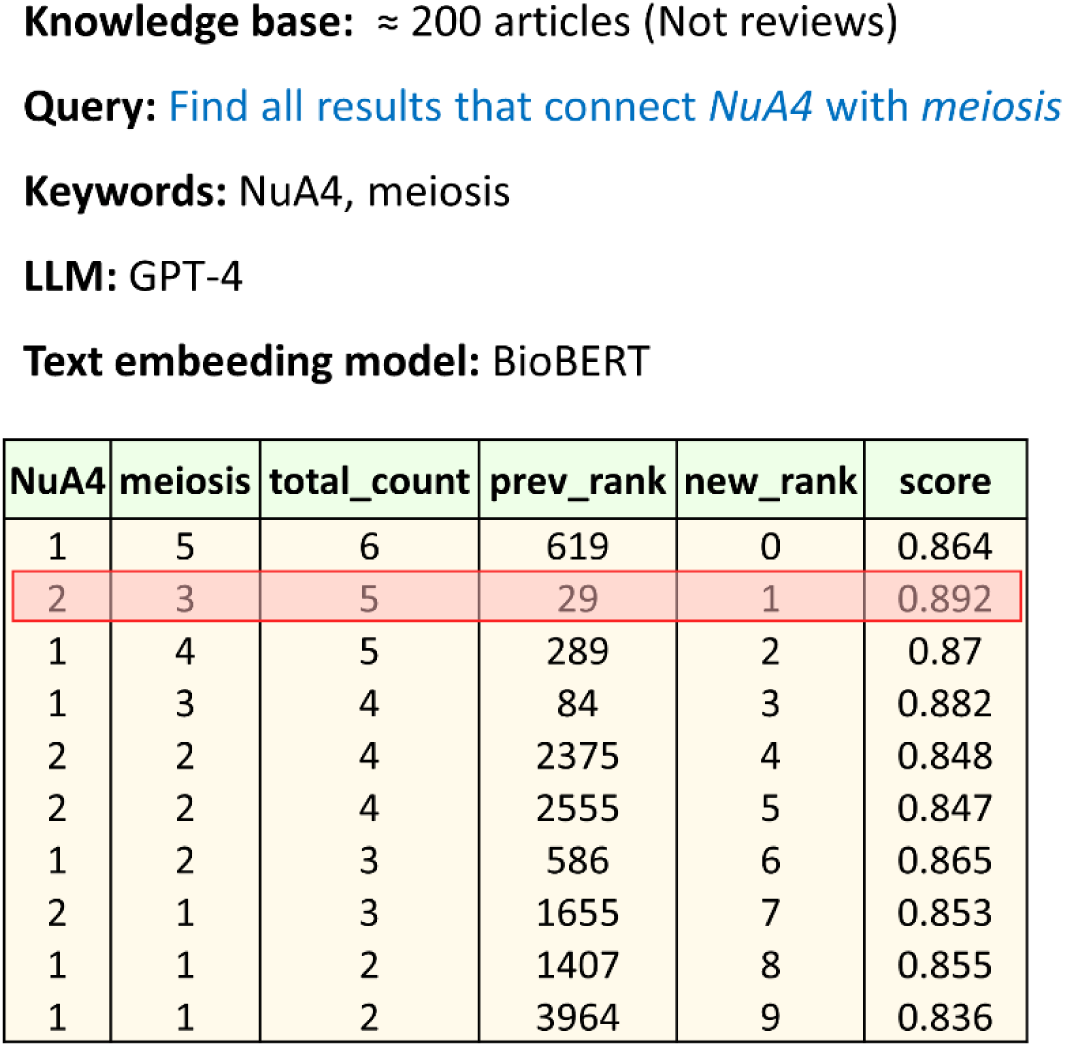
Ranking chunks by keyword frequencies ranks the key chunk higher –. The key chunk which was placed at 29^th^ place by PES1 approach is placed at second position after employing PES2.

**Table 2:** PES1 vs. PES2 based on LLM generated answers.

| Question | PES1 | PES2 |
| --- | --- | --- |
| What is known about viral chromatin? | IDK | Viral chromatin, the DNA-protein complex of viruses within host cells, undergoes epigenetic modifications like DNA methylation and histone changes, impacting latency and replication. It is dynamically remodeled by both cellular and virus-encoded proteins, crucial for gene expression regulation and latency establishment. |
| What is the role of r-loops at centromeres? | IDK | R-loops at centromeres play key roles in transcription, DNA repair, and chromosome segregation, affecting replication fork progression and centromeric repeat stability. They influence CENP-A abundance, recruit centromere proteins, and activate the ATR checkpoint pathway to ensure accurate chromosome segregation. |
| What is the evidence for phase separation in the nucleus? | IDK | Evidence for nuclear phase separation includes acetylated nucleosome-dependent phase separation, multi-bromodomain proteins inducing LLPS in acetylated chromatin, and chromatin phases formed by histone modifications. Additionally, proteins like TRNP1 and SAFB, RNA's role in MLO assembly, and starvation-induced proteasome assemblies demonstrate phase separation's impact on nuclear organization and function. |
| Is there any relation between E1A and p400? Elaborate on that | IDK | E1A, an adenoviral protein, interacts with p400, influencing apoptosis, transformation, and gene expression. This interaction is crucial for E1A's functions, as it stabilizes Myc by promoting Myc-p400 complex formation, essential for oncogenesis and the activation of Myc target genes. |
| Does TIP60 get modified upon DNA damage? | IDK | Yes, TIP60 gets modified upon DNA damage. |
| Can you find a link between TIP60 and r-loops? | IDK | Yes, the text indicates that TIP60–p400 binding to many target genes is enhanced by promoter-proximal R-loops, which are RNA:DNA hybrid structures. This suggests that R-loops play a role in the recruitment or stabilization of TIP60–p400 binding at these loci. |
| What is the functional relationship between the DCC and CLAMP? | IDK | CLAMP facilitates the DCC's binding to the male X chromosome's high-affinity sites in Drosophila, enhancing chromatin accessibility and transcription for dosage compensation. It acts as a pioneer factor, crucial for the coherent localization and function of the DCC. |
| Can you elaborate on the roles of MSL2 complex in | IDK | The MSL2 protein is key in Drosophila's MSL complex for dosage compensation, ensuring equal gene expression from the male X chromosome to match female levels. It forms a core complex with MSL1, enabling chromosome-wide |
| dosage compensation? |  | upregulation of genes via histone modification, specifically H4K16 acetylation. MSL2's expression is male-specific, regulated to prevent female dosage compensation. |
| Can you elaborate on the roles of SWR1 complex in yeast? | IDK | The SWR1 complex in yeast swaps H2A-H2B dimers with H2A.Z-H2B, affecting gene expression, chromosome segregation, and centromere silencing. It is essential for incorporating H2A.Z at gene promoters, influencing meiosis, and responding to environmental stress. |
| Find all results that connect NuA4 with meiosis | IDK | The NuA4 complex, especially through ESA1, affects meiosis by acetylating histone H4, influencing chromatin structure, homolog synapsis, and chromosome segregation. It plays a crucial role in gene expression regulation and chromatin remodeling, essential for spermatogenesis and meiotic efficiency. |

IDK → I do not know.

**Table 3:**
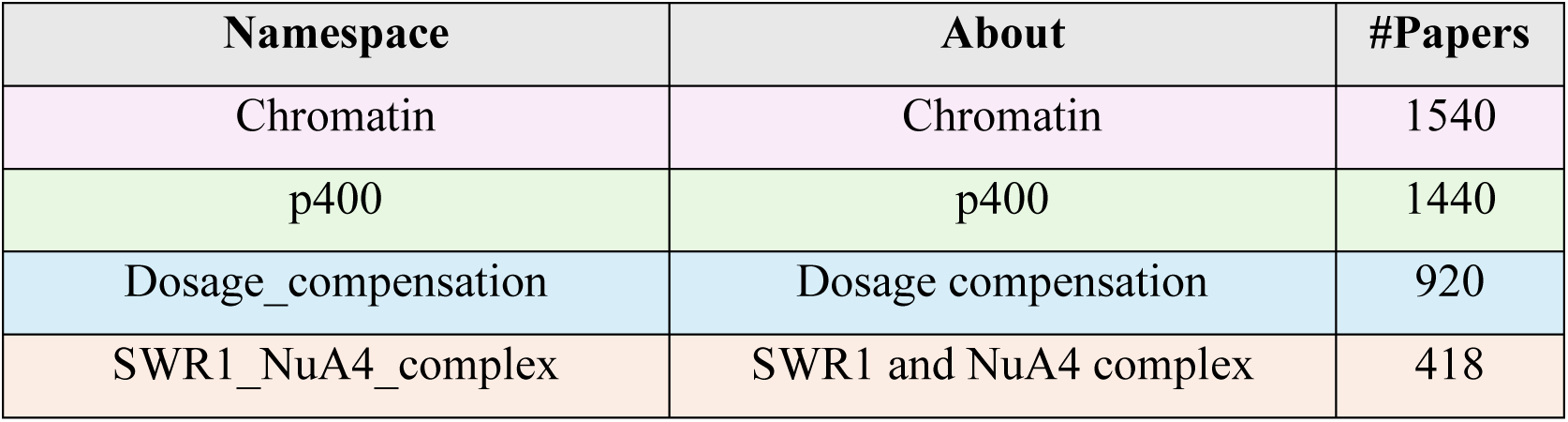
Namespaces with number of papers.

For both prompt enhancement approaches, our input prompt comprised of three pieces of information (see Fig 4A):

**a) Header:** This contains instructions that an LLM is supposed to follow.
**b) Context:** This is where the context to answer a question is placed.
**c) Question:** This is a placeholder for user questions.

As shown in Fig 4A, we explicitly instructed the LLM in the header not to make up an answer and seek it only within the provided context and if, and only if, an answer is not contained in the context then respond with ‘I don’t know’. It is clear from Table 2 that PES1 struggled to generate meaningful prompts as it responds with ‘I don’t know’ for all the questions. Furthermore, we evaluated the contexts produced by both PES1 and PES2, and we noticed that the information content in contexts extracted by PES1 is not relevant enough to answer questions listed in Table 2. This was even worse for questions that seek extremely specific details. On the other hand, with PES2 we were able to get answers to all of the questions. We provide all the detailed answers along with references in the in the supplemental Table 1.

## Discussion

### 1. Performance of PES1

Our experiment with different embedding models in PES1 highlights an interesting aspect of text embedding. The performance of BioBERT, BioGPT, and OpenAI’s ADA in ranking key chunks from biomedical literature can be influenced by a number of factors, including their design, training data, and algorithms.

#### BioBERT’s Performance

BioBERT is pretrained on a large corpus of biomedical texts which is likely to give it an edge in understanding and contextualizing biomedical queries. However, it ranked the key chunk much lower (29^th^ place) in a bigger knowledge base than it did in a smaller setting (2^nd^ place). This suggests a limitation in its retrieval capabilities as the complexity of knowledge base increases. This indicates that as the size of the knowledge base grows, even a domain-specific model like BioBERT might struggle with the increased diversity and complexity of the data, leading to lower quality information retrieval.

#### BioGPT’s Performance

BioGPT, similar to BioBERT, is tailored for biomedical content but there is an important distinction: the focus of the BERT model is to understand the meaning of words in relation to the entire text, whereas GPT excels at text generation and completion. So, as a generative model, BioGPT approaches embedding of texts differently which in turn influences ranking.

#### OpenAI’s ADA Performance

Being the worst performer, OpenAI’s ADA highlights a broader issue with general-purpose embeddings in domain-specific tasks [29–31]. ADA is trained on variety of texts, not just biomedical literature. When compared to a model like BioBERT, this generalist approach may limit its ability to understand and prioritize the complexities of biomedical text. Therefore, the embeddings generated by ADA may not capture domain-specific terms and relationships as well as BioBERT or BioGPT, resulting in poorer performance in specialized tasks. ADA’s ranking of the key chunk beyond the reach (9^th^ place) of GPT-3 even for such small set of chunks emphasizes the challenge that general-purpose embeddings face in domains with a specific focus. The model’s training on broader corpus may make it more sensitive to the specific cues that indicate relevance in biomedical texts but doing that comes with additional expense in terms of expertise, time, and money.

### 2. Performance of PES2

By re-ranking text chunks based on the frequency of specific keywords extracted from user queries, PES2 prioritizes content that is apparently more relevant to the users’ informational needs. <u>Strengths of the Keyword Frequency-Based Re-Ranking</u>:

**a.** <u>Improved relevance</u>: The results show that our re-ranking strategy significantly improves the relevance of retrieved chunks compared to the embedding-based approaches which mainly rely on cosine similarity measures in vector space. PES2 more efficiently surfaces information that users are likely to find helpful by directly aligning the retrieval process with the explicit keywords in user queries.
**b.** <u>Simplicity and efficiency</u>: In contrast to intricate NLP methods that necessitate substantial computational power and advanced AI models, keyword frequency analysis is reasonably easy to apply and computationally effective. For applications where quick response times are essential or resources are scarce, this makes it especially appropriate.
**c.** Flexibility: The approach offers considerable flexibility, allowing for easy adjustment of the re-ranking algorithm to accommodate different keyword selection schemes or to incorporate more relevance signals as needed.

The PES2 approach represents a promising advancement toward the creation of more relevant and responsive biomedical question-answering systems. This method provides a practical trade-off between retrieval effectiveness and computational efficiency by emphasizing the explicit signals supplied by user queries. However, realizing its full potential will require addressing its current limitations:

**a.** <u>Handling of synonyms and related terms</u>: A drawback of the PES2 approach is its dependence on precise keyword matches, which might leave out domain specific nuance like synonyms and related terms. In biomedical texts one keyword may have multiple synonyms, for example, Heterochromatin protein 1, Hp1a and Su(var)205 all refer to the same entity.
**b.** <u>Typos and Spelling Variations</u>: Spelling errors can have a substantial impact on the retrieval process and may result in the exclusion of valuable information.
**c.** <u>Contextual relevance</u>: Although keyword frequency can be used as a stand-in for relevance, it does not take the context of a keyword’s appearance into consideration. As a result, this approach may give preference to sections that have a high keyword density but provide little insight into the user’s actual query.

To address these limitations, future work could investigate the integration of semantic analysis techniques to better understand the context and meaning behind user queries. To include synonyms and related terms in the search, one possibility could be exploring query expansion techniques using resources such as the Unified Medical Language System (UMLS) which can aid in broadening the search query for biomedical texts by including synonyms or related terms. This increases the likelihood of finding relevant chunks even in cases where the exact keywords are misspelled. Typos and slight variations in the user’s query can be explored by fuzzy matching algorithms. To locate close matches to the keywords in a database, methods such as Levenshtein distance (a type of edit distance) can be employed. This way, relevant chunks can still be retrieved based on the closest matches to the intended keywords, even in the event of a typo. To improve contextual relevance, future work could include the integration of knowledge graphs. The graph can be used to enable semantic search functions. Understanding the semantic relationships between entities can be done by looking at the graph’s structure rather than just depending on keyword frequencies. This can help in ranking chunks not just by the occurrence of keywords but by the relevance of the context in which those keywords appear.

## Conclusions

Our findings emphasizes the importance of integrating external knowledge with generative AI to improve biomedical QA tasks. Through our proof-of-principle study, we demonstrated the significance of prompt enhancement strategy based on explicit signals in user’s query over traditional text embedding based context retrieval approaches. We also highlighted the importance of namespaces in streamlining the search, enabling users to experiment with different chunk sizes for enhanced precision in answers. Our method presents a viable strategy to strike a balance between retrieval efficiency and computational demands. Moving forward, unlocking the full potential of this method depends on addressing its current constraints. However, the insights derived from this investigation hold promise for broadening the horizons of NLP research and its application in solving complex QA challenges, especially in specialized fields like biology and biomedicine.

## List of abbreviations

NLP: Natural Language Processing
LLMs: Large Language Models
QA: Question-Answering
DB: Database
APIs: Application Programming Interfaces
PESs: Prompt Enhancement Strategies
RAG: Retrieval-Augmented Generation
KB: Knowledge Base
BERT: Bidirectional Encoder Representations From Transformers
GPT: Generative Pre-trained Transformer
PES1: Context Retrieval Based on Vector Similarity Ranking
PES2: Context Retrieval Based on Keyword Frequency Ranking
UMLS: Unified Medical Language System

## Declarations

### Ethics approval and consent to participate

Not applicable.

### Consent for publication

Not applicable.

### Availability of data and materials

WeiseEule is implemented as a localhost web application that can be accessed at the following GitHub repository: https://github.com/wasimaftab/WeiseEule-LocalHost Step-by-step instructions to make the app run on a user computer are also provided in the above link.

### Authors’ contributions

WA and TS conceived the study. WA designed and conducted the experiments, developed the application, and wrote the manuscript. TS and ZA contributed by providing several of the queries presented in Table 2, evaluated the application, and offered valuable feedback to refine both the application and the manuscript. KB inspired the conception of this work and provided invaluable feedback on the manuscript. All authors have thoroughly read and approved the final manuscript.

## Acknowledgements

The authors thank Prof. Peter B Becker from Molecular Biology Division of Biomedical Center, LMU Munich, Germany for providing valuable feedback on the manuscript.

